# A hybrid CNN-Random Forest algorithm for bacterial spore segmentation and classification in TEM images

**DOI:** 10.1101/2023.04.03.535316

**Authors:** Saqib Qamar, Rasmus Öberg, Dmitry Malyshev, Magnus Andersson

## Abstract

We present a new approach to segment and classify bacterial spore layers from Transmission Electron Microscopy (TEM) images using a hybrid Convolutional Neural Network (CNN) and Random Forest (RF) classifier algorithm. This approach utilizes deep learning, with the CNN extracting features from images, and the RF classifier using those features for classification. The proposed model achieved 73% accuracy, 64% precision, 46% sensitivity, and 47% F1-score with test data. Compared to other classifiers such as AdaBoost, XGBoost, and SVM, our proposed model demonstrates greater robustness and higher generalization ability for non-linear segmentation. Our model is also able to identify spores with a damaged core as verified using TEMs of chemically exposed spores. Therefore, the proposed method will be valuable for identifying and characterizing spore features in TEM images, reducing labor-intensive work as well as human bias.

## Introduction

Bacterial spores, also known as endospores, are dormant forms of sporulating bacteria that exhibit no cellular activity [1]. Spores are exceptionally resilient to external stressors such as temperature, humidity, radiation, and chemical exposure [2]. Due to their inherent resilience and ability to germinate back into bacteria when returned to more favorable conditions, spores from pathogenic bacteria pose a significant problem in many areas of society, including healthcare, food production, and homeland security [3–5]. Therefore, studying spores are important for developing new sterilization and detection strategies. To study spores and determine morphology, size, ultrastructure, topography, and structural features, Transmission Electron Microscopy (TEM) can provide valuable information. In particular, TEM enables high-resolution visualization of all the layers within a spore, which for example can provide important clues of the mode of action of light or disinfection chemicals [6, 7].

Spores are complex structures made up of several layers, including the core, cortex, coat, interspace, and exosporium [8]. Some species also express surface filaments as a “fluffy” layer or long fibers [9], as shown in Fig 1. These layers are all important components of bacterial spores and they have separate functions. The core is located in the center of the spore and contains the bacterium’s genetic material and cellular machinery, surrounded by protective chemicals such as dipicolinic acid (DPA). In addition, the core is covered by a membrane and cell wall that form the outer layers of the spore. The cortex, made up of peptidoglycan, surrounds the core, maintains the shape of the spore, and provides the initial energy source for the spore during germination. The spore coat, composed primarily of tightly packed protein layers, further surrounds the cortex. Finally, the interspace is mostly empty surrounding the coat, and is delimited by the thin exosporium layer consisting of proteins and lipopolysaccharides [10].

**Fig 1.**
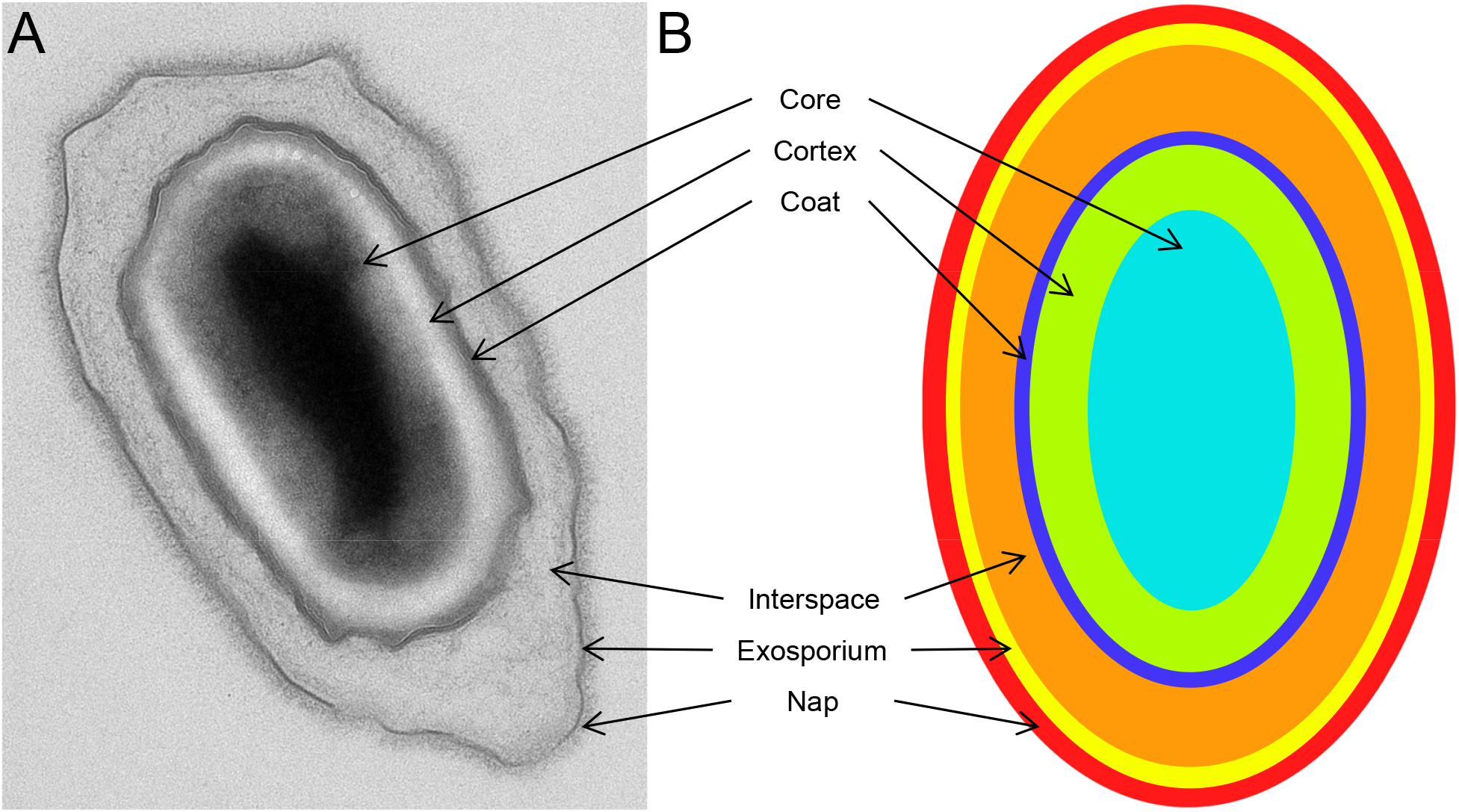
TEM micrograph (A) and structure model (B) showing the layer structure of bacterial spores. From the central spore core, the layers are in ascending order; cortex, coat, interspace, exosporium, and nap. The nap is the fluffy layer consisting of thin surface polymers (pili).

When spores are exposed to various decontamination agents, for example, sodium hypochlorite and peracetic acid, these agents change the spore’s structural composition. These changes the chemical and layer integrity of the spore, as observed using TEM [11]. Although useful in describing qualitative changes in the spore layer structure, these TEM observations are unsuitable for large quantitative evaluations since it is a very time-consuming process to analyze many spores. The analysis is, in addition, prone to human error by its nature.

One way to efficiently analyze a large number of TEM images, and avoid human bias during the assessment, is by using computerized methods like machine learning (ML) and deep learning (DL). ML is an automatic tool requiring little human input that can be trained to automate spore segmentation. ML methods have been applied in a variety of fields, such as healthcare [12], speech recognition [13], agriculture [14], and business forecasting [15], to improve the efficiency of a wide variety of processes. In particular, DL, a subset of ML, has been successfully applied in areas such as object segmentation, classification [16, 17], image recognition [18], autonomous vehicles [19], pattern recognition [20], etc. However, DL requires a very large amount of data to create reliable models and is, as such, limited in its applicability.

Over the years, ML methods have been developed to automate the identification and classification of microorganisms based on microscopic images. Li and colleagues [21] provided a comprehensive review of various analysis methods for content-based microscopic image analysis (CBMIA), including pre-processing, feature extraction, post-processing, and classification. Similarly, Kulwa and colleagues evaluated image processing and ML methods designed for segmenting microorganisms in images [22], while Li and colleagues reviewed clustering methods for analyzing microorganism images [23]. Additionally, Vikrant and colleagues presented an approach for microscopic image classification that combined guided image filtering, Otsu thresholding, and scale-invariant feature transform [24]. Ma and colleagues conducted a review of microorganism image analysis, exploring traditional image processing and traditional ML as well as DL methods for microorganism detection [25]. These related works primarily focus on ML approaches for microbiological image recognition, using either traditional ML or DL. While ML works well with limited data, DL requires a substantial amount of data but provides excellent performance. As a result, using a hybrid learning method that combines both traditional ML and DL approaches has the potential to achieve the benefits of both.

In this work, we develop an automated algorithm for segmenting and classifying layers in spore TEM images. The proposed algorithm combines a CNN for feature extraction and an RF classifier for pixel classification. We train the CNN with image data and use its predictions as input features to decision trees in the RF algorithm. Thus, the CNN converts high-dimension 2D TEM images into low-dimension features that preserve the locality of pixels and reduce the curse of dimensionality for accurate prediction through the RF classifier. Evaluation of the proposed CNN-RF algorithm shows that it performs better than other state-of-the-art algorithms and can both accurately and efficiently analyze large amounts of spores in TEM images.

## Theoretical background and definitions

### Convolutional Neural Networks

Convolutional Neural Network, introduced in 1989, is a method inspired by how the human visual cortex in the brain processes visual inputs into information [26]. In a CNN, convolutional layers are the fundamental building blocks [27]. A convolutional layer performs a set of convolution operations on the input data, which are a combination of element-wise multiplications and summations. The convolution operation is performed between the input data and a set of filters, also known as kernels or weights, that are learned in the training process. The filters slide across the input, computing a dot product between the filter and the input data [28]. The number of filters determines the number of feature maps that are generated by the convolutional layer, each capturing different features or aspects of the input data, which for this work are 2D TEM images.

Another important component of CNNs is pooling layers. These down-samples the input data and reduce its spatial dimensions [29]. Downsampling is useful for two reasons, to reduce the computational cost of the network, as the amount of data to be processed is reduced, and to make the representations learned by the network more invariant to small translations of the input data. This thereby controls overfitting by reducing the number of parameters. Overfitting occurs when a model starts to memorize the characteristics of the training data and, in turn, loses its ability to generalize. To also increase non-linearity in the network an activation function can be used for the feature maps. As activation function, the rectified linear unit (ReLU) is commonly used, which is then applied to the output of each neuron in the network to learn a wider range of complex representations and to improve the ability to classify images [30]. The ReLU function returns the same input for positive values and returns 0 for all negative [31].

### Decision Tree

In applications where the aim is to classify items into classes, decision tree algorithms are often used. It works by building a tree-like model of decisions based on feature values. The tree is constructed and uses an algorithmic approach that searches for features that group the data more homogeneously [32]. A decision tree thereby predicts the class label of each pixel in the image. To determine how the features should be optimally split into nodes in the tree, the Gini impurity measure can be used. The Gini impurity thus measures misclassification of randomly drawn samples from each node. The Gini impurity decreases as nodes are added to the tree and when the Gini impurity is zero, the node is not expanded. The Gini impurity of a node *n* is calculated as,

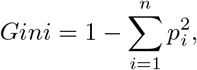

where *n* is the number of classes and *p*_*i*_ is the probability of node *n* in class *i*. In a decision tree, an input sample is thereby checked against each of the conditions at each node, and a node’s offspring is selected depending on whether the condition is True or False.

### Random Forest

Random forest is an ensemble approach that contains multiple decision trees to make predictions from data [33, 34]. The decision trees in a RF are trained using a process called bootstrapping, which involves sampling the data with replacement. This implies that some data points may be included in the training set more than once, while others may not be included at all. Each decision tree in the RF makes predictions based on the features in the data, and the majority voting calculates the final prediction, which is based on individual decision trees’ output. By using multiple decision trees, the risk of overfitting is reduced compared to using a single decision tree. Thus, each tree randomly selects a subset of features from the whole data set. With this method, training results are obtained based on different feature sets, and sampling with return ensures that training results are valid and reliable.

### Design of the CNN-RF algorithm

We design our algorithm as a CNN-RF algorithm to optimally segment spores and classify layers. The CNN is first trained on image data to convert high-dimensional 2D TEM images into vectors of real values, which are then used as input features for decision trees in the RF algorithm. By preserving the inter-pixel relationships in the image, the features extracted from the CNN architecture enhance the accuracy of the prediction, while also reducing the dimensionality of the features [35]. The CNN thereby serves as the feature generation step, which is followed by the RF classifier for precise classification.

### Spore preparation and TEM image acquisition

*Bacillus thuringiensis* ATCC 35646 cells were grown on BBLK agar (210912, BD) plates and set to incubate at 30°C overnight. These cells were collected by scraping them off the agar and transferring them to a 1.5 ml Eppendorf tube, after which they were centrifuged to remove leftover growth media. To allow sporulation, the cells were stored at 4°C overnight. Before use, the sporulated suspension was rinsed five times by centrifuging in deionised water for 5 minutes at 5000 *G*.

To prepare spores for TEM the suspensions were fixed with 2.5 % Glutaraldehyde (TAAB Laboratories, Aldermaston, England) in 0.1 M PHEM buffer and further postfixed in 1 % aqueous osmium tetroxide. The spores were then further dehydrated in ethanol, acetone and finally embedded in Spurr’s resin (TAAB Laboratories, Aldermaston, England). 70 nm ultrathin sections were then post contrasted in uranyl acetate and Reynolds lead citrate. Spores were imaged using a Talos L120C (FEI, Eindhoven, The Netherlands) operating at 120kV. Micrographs were acquired with a Ceta 16M CCD camera (FEI, Eindhoven, The Netherlands) using TEM Image and Analysis software ver. 4.17 (FEI, Eindhoven, The Netherlands).

### Annotation of TEM images

Accurate and detailed annotation of TEM images provides important context for understanding the features and regions within the image, and serves as the ground truth for ML training. We labeled eight distinct categories within a TEM image of a spore, “coat”, “core”, “cortex”, “exosporium”, “interspace”, “nap”, and “background”. For areas of the image that were not part of the spore or background, like debris, and for areas of the spore that were smeared or otherwise could not be resolved due to poor sectioning or overlapping areas, we used the label “bad region”. We used APEER (APEER by Zeiss, 2022) for annotation [36] to easily and efficiently label the TEM images with the different categories. This web client software provides a user-friendly interface with tools for creating labels, selecting the appropriate category for each region, and saving the annotations in a format compatible with ML algorithms. This ensures that the annotations are accurate, reproducible, and can be used for future analysis.

### Data preprocessing, training and testing data

Before feeding the TEM image data into the model, we employed some necessary steps to ensure better model performance. First, we resized all training images from 3000×2500 to 2048×1664 pixels and normalized the data between 0 and 1. This resizing and normalization process was essential for improving the model’s performance, as it ensured that all the extracted features had the same value range. Second, we used a data augmentation technique to expand the number of images in the data set. By applying various image transformations, such as rotation, scaling, and flipping, to the images in the training set, we were able to increase the diversity of the training data and reduce the risk of overfitting. This augmentation technique proved highly effective, as it helped the model learn more robust features that generalize well to new images. We augmented 64 images from the training data to create 384 new images. During training, we passed augmented data to the CNN to extract abstract features from the data set, and these features were sent to the RF classifier. The CNN model used a series of convolutional layers to learn the relevant features from the input images, while the RF classifier was used to classify the spore classes. After training a model on a set of images, it is crucial to evaluate its performance on unseen data to determine its efficacy in real-world scenarios. To achieve this, we created a testing data set comprising 50 images that were distinct from the training data set. The trained model was then applied to this testing data, and its performance was evaluated by comparing the predicted spore class against the actual spore class. This allowed us to determine the accuracy and reliability of the model in predicting spore classes for previously unseen data. The results obtained from this evaluation helped fine-tune the model further to improve its performance. Overall, the approach employed during the model training helped in creating a robust and reliable model for predicting spore classes from images.

The model was trained on a computer with an Intel Core i9 processor, 32 GB of RAM, and an NVIDIA GeForce GTX 1600 SUPER graphics card. The training was performed using Python programming language, with the TensorFlow and Keras libraries [37, 38] for DL and scikit-learn library [39] for RF classifier. The total time taken to train the model was approximately 8 hours. This duration includes the time taken for data preprocessing, model training, and evaluation. During this time, the model was trained on a total of 384 images. The training was performed using the n_estimator value of 300 and a single decision tree was built with 25 features in RF.

### Model Architecture and data description

Our proposed architecture for spore segmentation is illustrated in Fig 2. The source code for the model implementation is available for access and download [40]. A step-by-step guide on how to install packages and run the model is provided in the supporting information. The CNN in our architecture employs 15 convolutional layers with ReLU layers and 5 max-pooling layers. The convolutional layers generate low-level features at the beginning and high-level features towards the end of the architecture, while the max-pooling layers help reduce the dimensionality of the extracted convolutional features. By utilizing a large number of convolutional layers, rectified linear unit layers, and max-pooling layers, the CNN can generate a high-dimensional feature space, and the RF classifier combines these features to make the final decision. The input image size used is 2048×1664×3, with 32 kernels of size 3×3 and stride 1 applied to each input image. The resulting 32 output feature maps are passed through the first block, which generates 64 features. Subsequently, each consecutive block generates 32 × 2^*n*^ features, where n = 2, 3, 4, or 5. Finally, the CNN predicts 1024 features of size 128×104.

**Fig 2.**
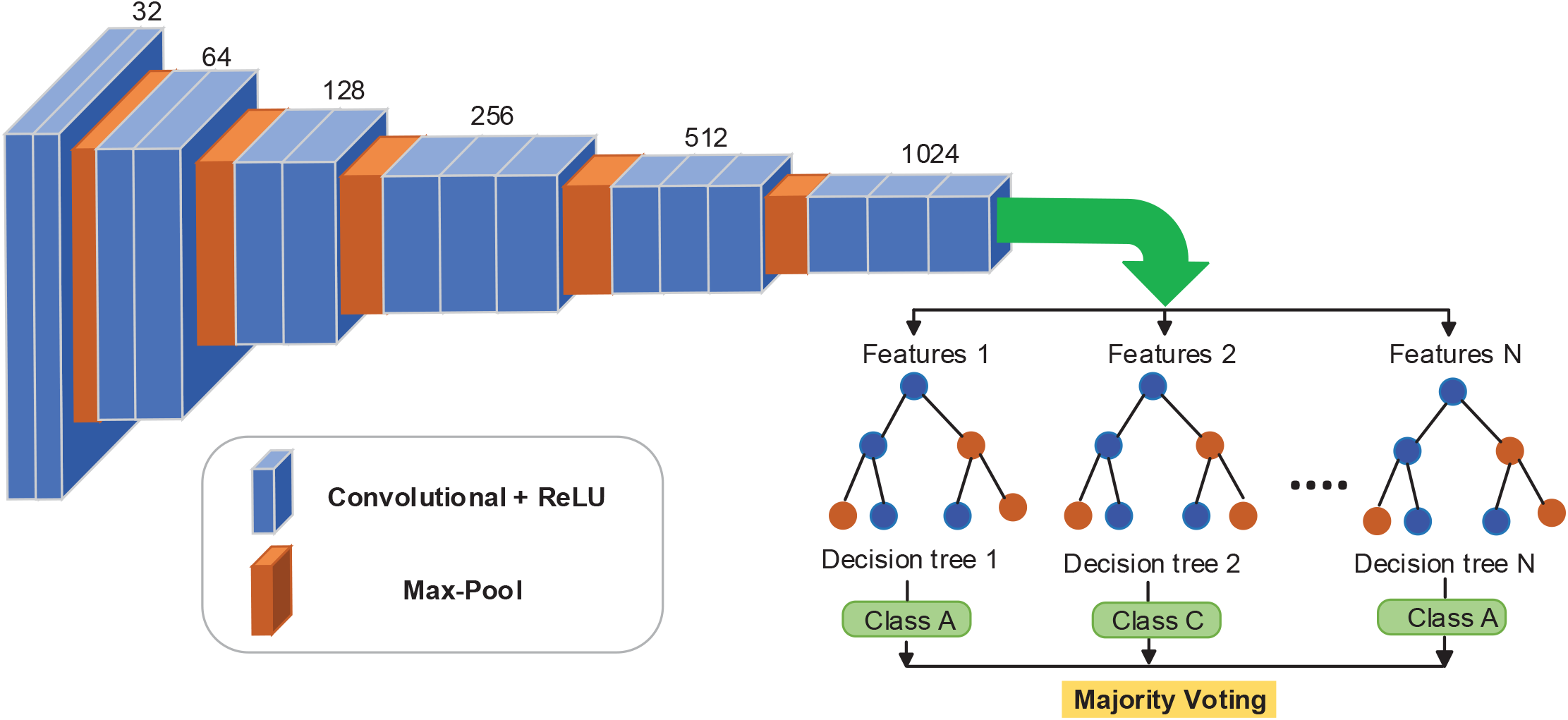
Proposed combined approach of CNN and RF. The CNN extract features from the data and RF classifies data based on a large number of decision trees.

During training, the model uses a three-fold cross-validation approach, where each fold *k* = 1, 2, 3 uses two-thirds of the whole data set for training. The remaining data is used as the test set. In each iteration, the CNN is trained on the sub-training data (*t*_*k*_ – *t*_*ki*_) and validated on the subset *t*_*ki*_. This approach enables the model to learn from different variations of the data and ensures that the model generalizes well to unseen data.

Once the CNN is trained, the decision tree predicts the spore classes based on the CNN features. To improve the robustness of the model and reduce the impact of errors made by individual decision trees, the majority voting approach is used to make the final prediction using the results from all decision trees. Finally, the performance of the trained model is evaluated on the test set to measure the model’s performance on unseen data.

#### Algorithm 1

Proposed Approach Pseudo code

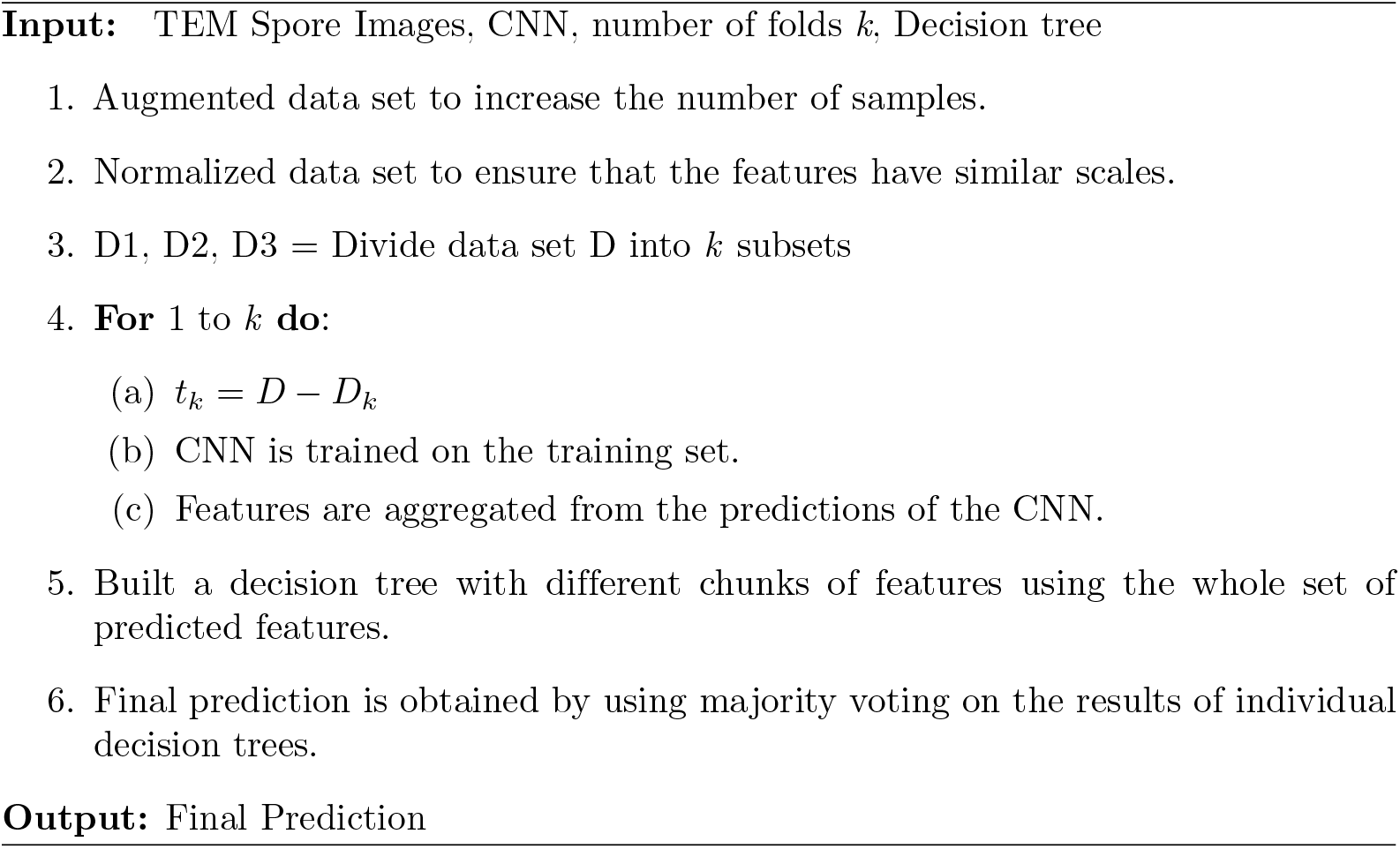

The proposed method is presented in Algorithm 1 in pseudo-code form. To increase the data sample and ensure that the features are on the same scale, augmentation and normalization are applied in lines 1-2. The three-fold validation process begins by dividing the data into three subsets. In line 4, these subsets are used to train the CNN, which then predicts features and aggregates them into numerical vectors. In line 5, decision trees are built using different chunks of features. The majority voting approach is used to obtain the final prediction. Therefore, the algorithm follows a straightforward process of data preprocessing, three-fold validation, training the CNN, building decision trees, and making predictions using the majority voting approach.

### Statistical metrics used for evaluation

In ML, evaluating the performance of a classification model is essential. Therefore, it is important to choose the right evaluation metric for the specific problem being solved because it significantly affects the results. We used four metrics to evaluate the performance of the proposed model: accuracy, precision, sensitivity, and F1-score. These metrics are commonly used for image classification and segmentation tasks, and they are calculated based on the number of true positive (TP), true negative (TN), false positive (FP), as well as false negative (FN) predictions made by the model [41].

Accuracy is the number of correct predictions made by the model divided by the total number of predictions. It is a simple and straightforward metric that provides a general understanding of how well the model is performing. However, it can be misleading in cases where the data is imbalanced, implying that one class has significantly more samples than the other. In these cases, a model that always predicts the majority class will have high accuracy, even though it is not making any useful predictions for the minority class. The accuracy is defined as,

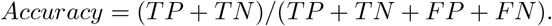

Precision is the number of TP predictions divided by the sum of the number of TP and FP predictions. Precision is a measure of how many of the positive predictions made by the model are actually correct. A high precision indicates that the model is not making many FP predictions, but it does not tell us anything about the FN predictions. The precision is defined as,

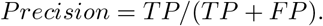

Sensitivity is the number of TP predictions divided by the number of TP and FN predictions. Sensitivity is a measure of how many of the actual positive samples are correctly identified by the model. A high sensitivity indicates that the model is not making many FN predictions, but it does not tell us anything regarding the FP predictions. The sensitivity is defined as,

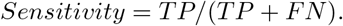

The F1-score is the harmonic mean of precision and sensitivity. It is a good metric to use when the data is imbalanced, as it takes into account both precision and sensitivity. The F1-score provides a balanced view of the model’s performance, as it considers both the FP and FN predictions. The F1-score is defined as,

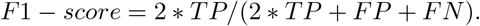

The final statistical metric is the support value in the confusion matrix. The support values represent the number of samples in the data set that belong to each class. It is the total number of instances that belong to a particular class and is often listed along the diagonal of the confusion matrix. The support value is important because it provides information about the distribution of the classes in the data set, which helps in evaluating the performance of a classification model more accurately. For instance, accuracy can be misleading if the data set is imbalanced. The support value helps to address this issue and allows for a more nuanced evaluation of the model’s performance, taking into account the distribution of the classes in the data set.

### Comparative statistics for chemically treated spores

The different spore samples treated with hypochlorite and with peracetic acid were compared to the untreated spores in their relative spore content (core, cortex, coat). We used Prism 9.3 (GraphPad Software) for statistical evaluation. The samples were compared overall using ANOVA and individual comparisons to respective controls were done using Dunn’s multiple comparisons tests.

## Results

To evaluate the proposed model for spore segmentation in TEM images, we employed a variety of metrics such as accuracy, precision, sensitivity, and F1-score as defined in the previous section. These metrics provide an effective means of assessing the performance of the model from different perspectives. For example, precision serves as a measure of the classifier’s exactness, and a low precision suggests a high number of FP. In contrast, sensitivity serves as a measure of the classifier’s completeness, where a low sensitivity implies a high number of FN. The F1-score considers both precision and recall (memory) and is considered to be most accurate when it is equal to 1, and least accurate when equal to 0. Furthermore, we used the trained model to classify individual images from the training and testing data sets and then calculated the accuracy of the predictions. To do this, we passed each image as input into the model, in which the model provided outputs as a prediction for the class of the image. After that, we compared this prediction to the true class of the image. We stored the accuracy of one-by-one images in an array and found the average performance on training and testing data sets. It gave an overall estimate of how well the model was able to classify the images in the data set. By finding the average accuracy, we also got a sense of how well the model was able to generalize to new, unseen images. Additionally, this approach also allowed us to inspect the model’s performance on specific images, and identify any images that the model may have struggled with.

### Classification accuracy assessment using a confusion matrix

We classified spore bodies into eight distinct categories: “Badreg”, “coat”, “core”, “cortex”, “exosporium”, “interspace”, “nap”, and “background”. “Badreg” (bad region) is related to regions not part of a spore. To evaluate the accuracy of our model, we employed a confusion matrix, which provides a visual representation of the number of TP, FP, FN, and TN predictions for each class, see Fig 3. The confusion matrix shows metrics for the proposed method based on test data. Fig 3(A) represents the number of instances in each cell belonging to each class. The diagonal cell of the confusion matrix represents the correctly classified instances for a class. By using the confusion matrix we could identify the strengths and weaknesses of our proposed model.

**Fig 3.**
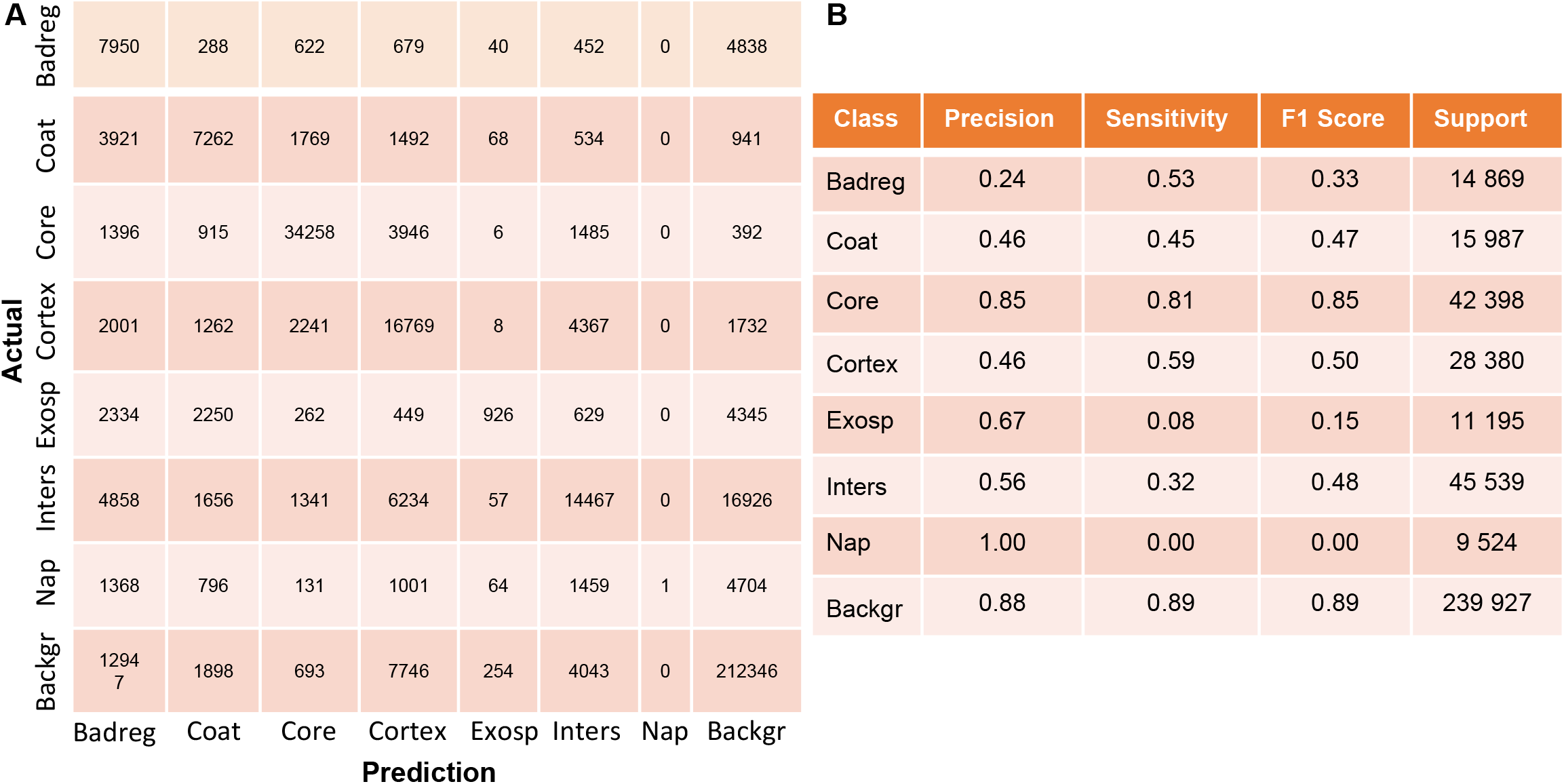
Classification performance for different spore layers based on the testing data set (A). Each cell shows the number of classified instances by the model, as compared to ground truth data. The diagonal cells show the correctly classified instances for a class. From this we evaluated the precision, sensitivity, and F1-score for each individual class (B).

Using the confusion matrix, we calculated the Precision, Sensitivity, F1-Score, and Support for each class in the data set, see Fig 3(B). Based on these metrics, we found that the proposed model performed well in predicting the different types of spore classes in terms of precision, sensitivity, and F1-score, particularly for “core,” “cortex,” and “background”. However, the model did not perform equally well when predicting the “exosporium” and “nap” classes. This likely originates from the close pixel values of the “interspace” class.

### Comparison with other classifiers

We used a RF classifier in our proposed model as the primary method for classification. To determine the effectiveness of the proposed RF classifier, we compared it with three other classifiers: AdaBoost (CNN-AdaBoost), XGBoost (CNN-XGBoost), and SVM (CNN-SVM). We evaluated the classifiers based on the same metrics as before. The results show that the proposed model, which used the RF classifier, achieved 100% accuracy, precision, sensitivity, and F1-score when trained on the data. This indicates that the RF classifier is capable of accurately classifying data with high consistency.

The second-best performer among the other classifiers was the CNN-SVM model, which achieved 81% accuracy, 86% precision, 56% sensitivity, and 58% F1-score. The support value of 965 693 was associated with different classes of training data.

During the testing phase, the proposed model using the RF classifier achieved 73% accuracy, 64% precision, 46% sensitivity, and 47% F1-score. These results indicate that the model performed well on the test data, although there was a decrease in performance compared to the training data. The second-best performer among the other classifiers during testing was the CNN-XGBoost model, which achieved 71% accuracy, 62% precision, 45% sensitivity, and 46% F1-score. The support value was 407 819.

Our experimental results suggest that the proposed model has stronger robustness and higher generalization ability compared to other classifiers for the non-linear problem of spore segmentation. The results of all classifiers are listed in Table 1.

**Table 1.**
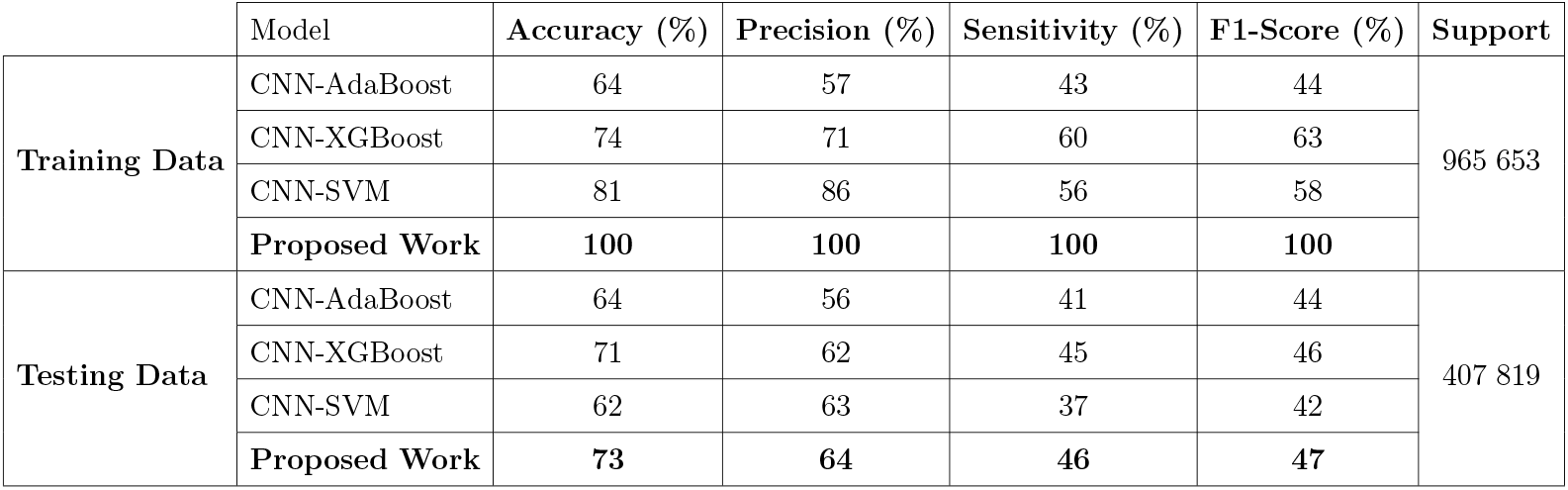
Comparing results for spore segmentation using different classifiers.

|  | Model | Accuracy (%) | Precision (%) | Sensitivity (%) | F1-Score (%) | Support |
| --- | --- | --- | --- | --- | --- | --- |
| Training Data | CNN-AdaBoost | 64 | 57 | 43 | 44 | 965 653 |
|  | CNN-XGBoost | 74 | 71 | 60 | 63 |  |
|  | CNN-SVM | 81 | 86 | 56 | 58 |  |
|  | Proposed Work | 100 | 100 | 100 | 100 |  |
| Testing Data | CNN-AdaBoost | 64 | 56 | 41 | 44 | 407 819 |
|  | CNN-XGBoost | 71 | 62 | 45 | 46 |  |
|  | CNN-SVM | 62 | 63 | 37 | 42 |  |
|  | Proposed Work | 73 | 64 | 46 | 47 |  |

### Accuracy during training and testing

In Fig 4, a histogram illustrates the distribution of the model’s accuracy across individual images in both the training and testing data to identify areas for improvement. The x-axis represents the range of accuracy values from 0-100%, while the y-axis represents the number of images in each accuracy range. The histogram indicates that the model achieves an average accuracy of 95.6% on the training images and 73.7% on the testing data. Most of the images in both data sets have accuracy values that fall within a narrow range, indicating the model’s consistent performance. The high average accuracy on the training data suggests that the model has learned to segment spores accurately. Nevertheless, the testing data shows a slightly lower average accuracy, suggesting that there may be opportunities for enhancement and additional optimization.

**Fig 4.**
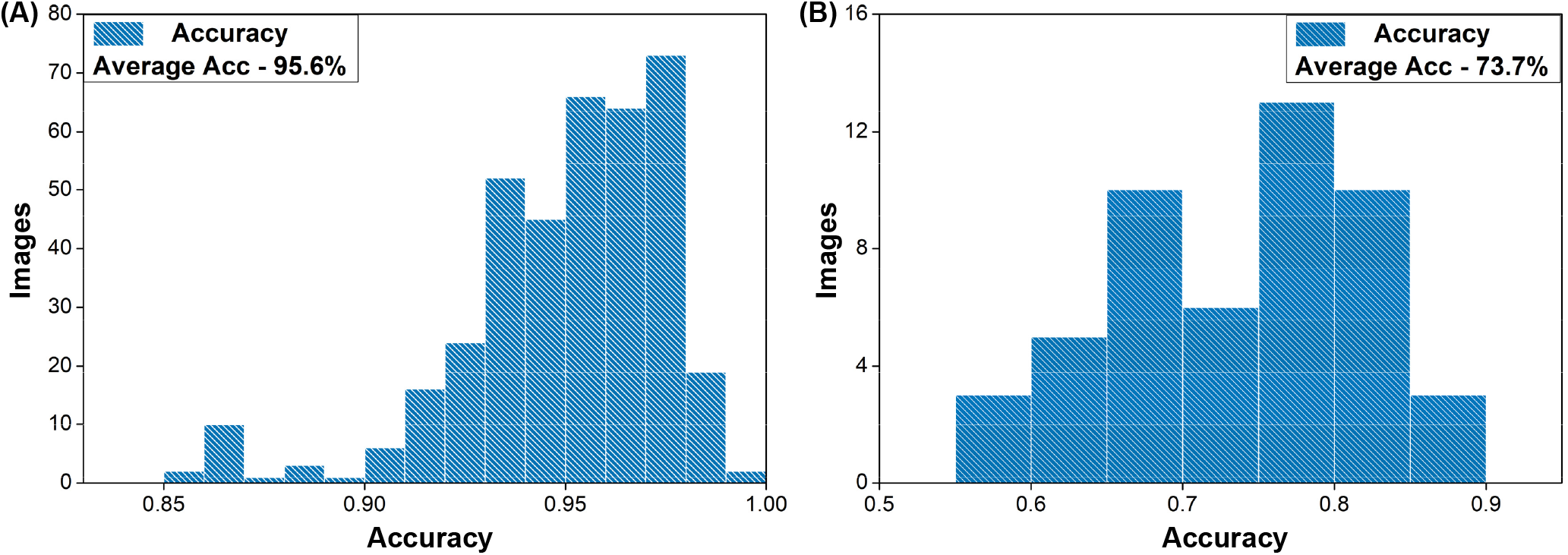
Distribution of the accuracy for the model when analysing all the individual images. (A) shows the distribution for the training data and (B) the testing data.

### Quantification of spore layers

Examples of TEMs, CNN feature extraction, and RF prediction are shown in Fig 5. The top 8 features at layers 4 and 10 are shown in supporting Fig S1. As mentioned in the method section, the CNN extract features from TEMs that represent spore layers, edges, and shapes by employing a series of convolutional layers. The dimensionality of the features is reduced using pooling, and the outcome of this process is a set of high-level features that represent the image. After this, the RF algorithm, uses these high-level features generated by the CNN to predict the segmentation of spore layers in the sample. The algorithm performs a pixel-wise classification of the image using the extracted features, and the output is a 128×104 matrix, with each entry representing a pixel in the spore sample. The value assigned to each entry in the matrix ranges from 1-8 and indicates whether a pixel is part of a spore layer. The individual classes’ segmented pixels can be seen on the right side of Fig 5.

**Fig 5.**
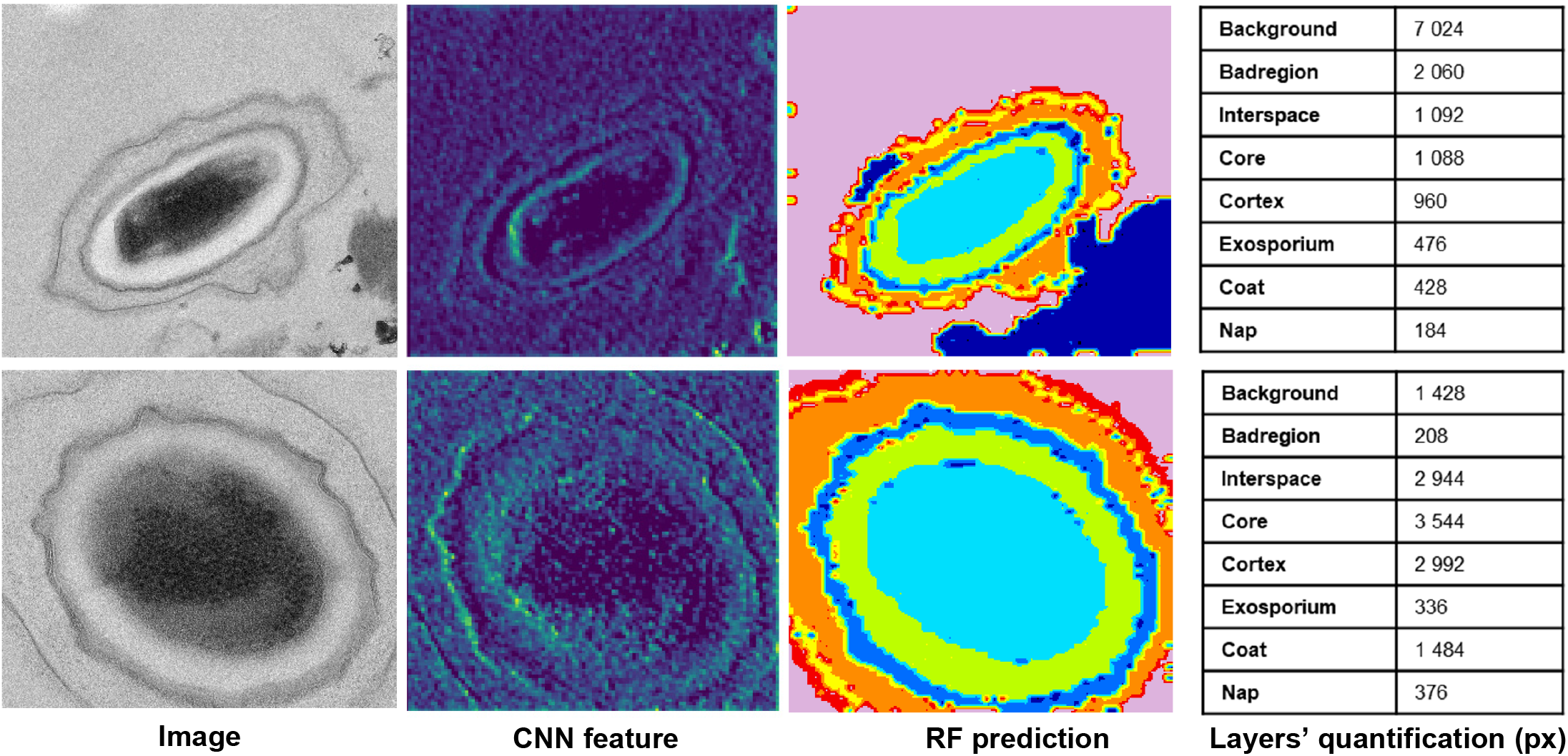
A sample of TEMs, CNN features, and RF predictions. The final segmentation map is a 128×104 matrix, with each entry representing a pixel in the spore sample. Right-side matrix shows segmentation for each individual class. The color coding for classification is “Badreg (dark blue),” “coat (blue),” “core (light blue),” “cortex (green),” “exosporium (yellow),” “interspace (orange),” “nap (red),” and “background (violet).”

### Visualize spore segmentation and classification in TEMs

We show in Fig 6 examples of spore segmentation and layer classification in TEM images using the proposed model. Original images are shown on the left, with manually labeled images in the middle. Ten more examples are shown in supporting Fig S2. On the right, the model’s prediction for the images is displayed with its corresponding accuracy. Note that the model can accurately identify the edges and boundaries of the objects in the image. As seen in the image, the model’s prediction closely aligns with the edges and boundaries of the labeled object, making it highly accurate in detecting objects within the image. Overall the results show that the model performed well and achieved an overall accuracy of 73% on the test data.

**Fig 6.**
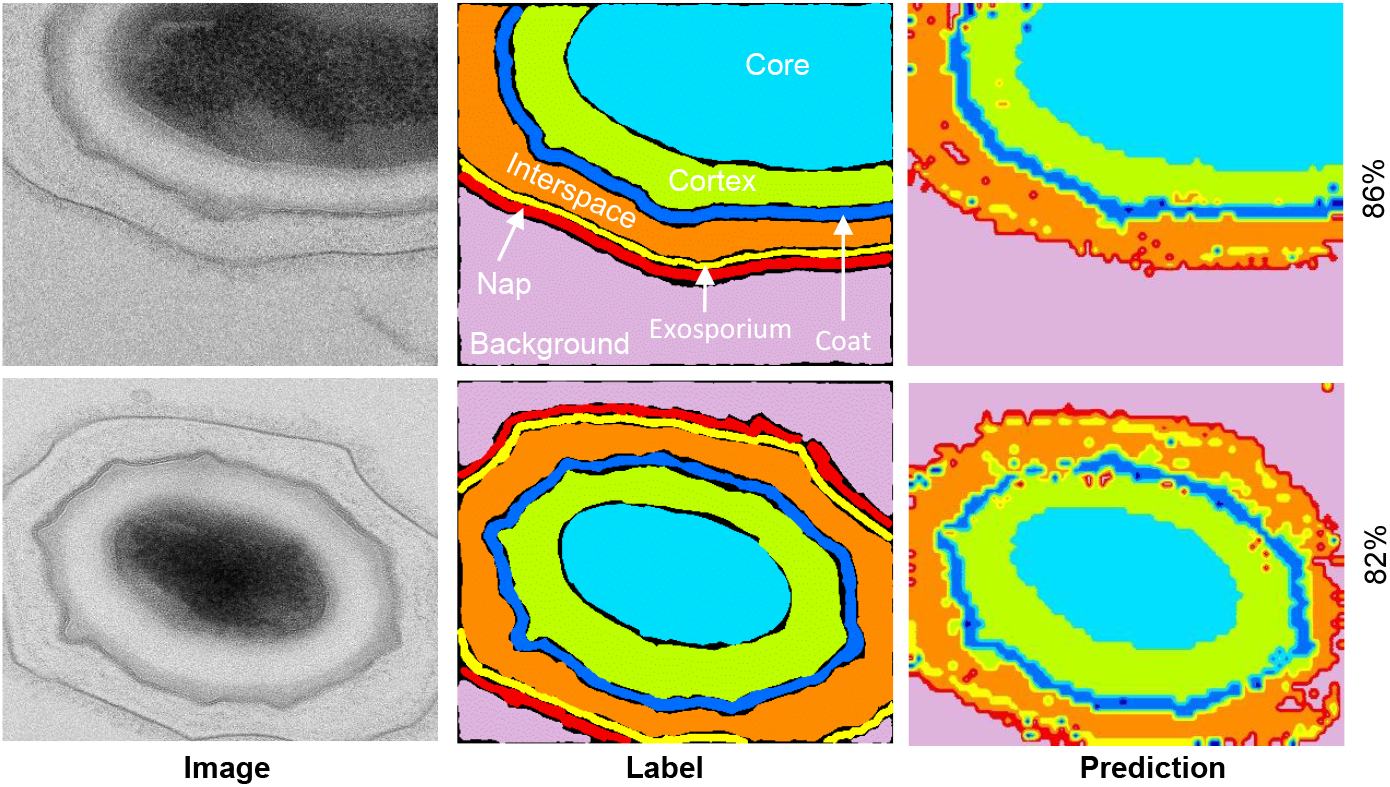
Comparative image showing TEM images of two spores, their respective layers as labeled, and their layers as predicted by our model.

### Analysing spore damage after chemical exposure

To also assess if the model could predict spore damage, we examined spores that were treated with a 0.5 % solution of sodium hypochlorite (commonly known as bleach) with a pH level of 11.55, and spores exposed to 1 % peracetic acid. These particular concentrations have been reported previously as being effective in killing spores [42, 43]. Sodium hypochlorite is a readily available and cost-effective decontaminant compound that acts by catalyzing several chemical reactions, such as saponification of fatty acids, neutralization, and chloramination of amino acids, thereby decomposing organic matter [44]. Hypochlorite-induced oxidative damage has been demonstrated to affect lipids, proteins, and DNA, as evidenced by previous studies using TEM [6]. For instance, the research showed that hypochlorite-treated spores underwent structural changes resulting in loss of integrity and discoloration of the core while exhibiting decomposition of the cortex, spore coat, and exosporium, ranging from defined structural traits to faint outlines with unstained content. On the other hand, peracetic acid is a type of disinfectant that can disable microorganisms through the oxidation of sulfhydryl and sulfur bonds, resulting in protein, enzyme, and metabolite denaturation [45]. However, unlike hypochlorite, its impact on spore integrity is less pronounced [6].

We analyzed TEM images to assess spore layer integrity after exposure to sodium hypochlorite and to peracetic acid. An example of a spore that lost the core integrity after sodium hypochlorite exposure is shown in Fig 7A, with the model’s pixel prediction in Fig 7B. This pixel-wise classification allows for a detailed and nuanced analysis of the spore layers. By breaking down the image into individual pixels and analyzing each one, the algorithm detects and classifies different types of spore classes based on their specific characteristics and features.

**Fig 7.**
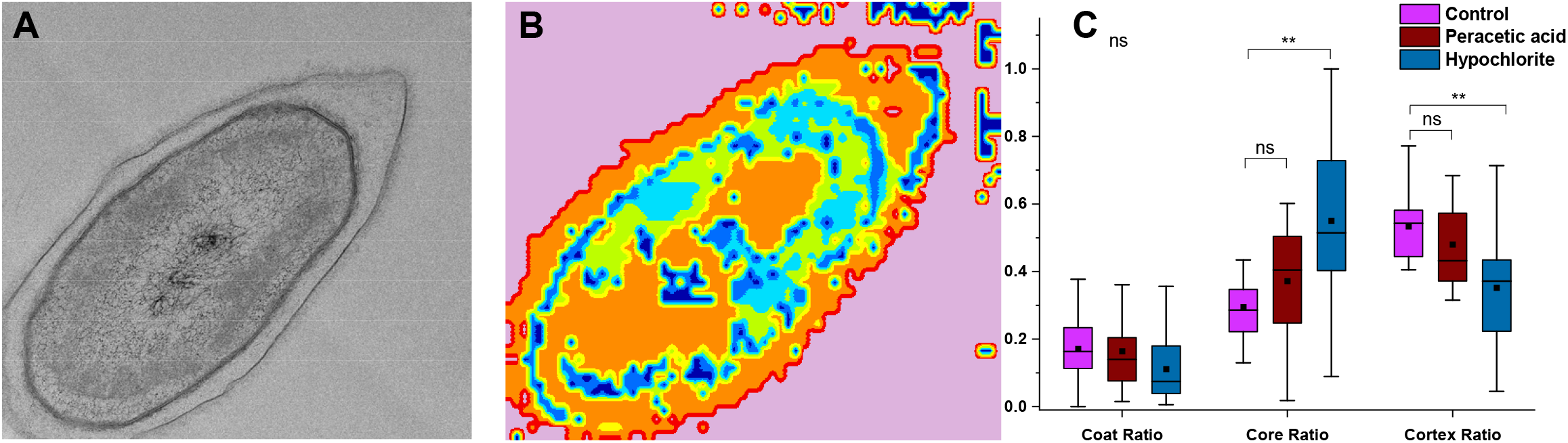
Assessing the spore layers integrity of sodium hypochlorite and peracetic acid exposed spores. A TEM image of a sodium hypochlorite exposed spore is shown in (A). The model’s classification is shown in (B). Coat, core, and cortex ratio for control spores, and hypochlorite as well as peracetic acid exposed spores. (C) shows the relative areas of the spore coat (n=33), core (n=22), and cortex (n=21). There was no significant difference in the coat ratio across the samples (indicated with “ns”), however, the hypochlorite-treated samples showed a significantly different core ratio (p=0.0014) and cortex ratio (p=0.0095).

We found that the algorithm correctly predicts most of the regions as background class, as in the original image. However, since the spores are damaged, the algorithm also predicts this damage. This is a significant finding since the algorithm was trained using a dataset that did not include images of sodium hypochlorite or peracetic acid. We also quantified the number of pixels classified as coat, core, and cortex for control spores (unexposed), and chemically exposed spores, Fig 7C. The model clearly identifies that the core integrity of chemically exposed spores is damaged by significantly overestimating the core ratio in comparison to the control. In addition, this overestimation reduces the cortex ratio for hypochlorite exposed spores. Thus, by assessing these two specific spore layers, it is possible to use our model to predict if the core of spores has been damaged by a chemical agent. Thus, we conclude that the algorithm is able to accurately analyze if the spore core has been damaged, indicating that our model has the potential to be used in real-world applications.

## Discussion

TEM imaging is a powerful approach when assessing the features of micron-scaled objects including bacterial spores. Nevertheless, this approach can be time-consuming and susceptible to human bias, especially when dealing with low-contrast images. To address these challenges, we develop in this work a CNN-RF algorithm optimized for segmenting and classifying spore layers in TEM images. To evaluate the performance of our algorithm, we conducted a comparative analysis against some commonly used classification algorithms, that is, Adaboost, Xgboost, and SVM. These methods have proven successful in various applications. Adaboost is a technique that combines multiple weak classifiers to create a more robust classifier. XGBoost is a gradient-boosting algorithm that is particularly effective for analyzing structured data, and SVM is designed to find the optimal hyperplane that maximally separates the different classes of image data in a high-dimensional feature space.

To achieve good performance our model utilizes a CNN to extract 1024 features from a single image while preserving the pixel locality, ensuring accurate prediction. Conversely, feeding TEM spore images directly at the pixel level violates this locality and leads to the curse of dimensions, which can have a negative impact on algorithm performance. However, the proposed RF performed well in their presence, whereas Adaboost, Xgboost, and SVM algorithms struggled when faced with irrelevant features in the data. The assessment of all methods on a test data set shows that our proposed model was better in all compared metrics.

To reduce the risk of overfitting and to handle imbalanced data, our method combines the strengths of both CNN and RF. The CNN handles imbalanced data by learning features that capture relevant patterns in the data. In contrast, RF provides a robust and accurate method for classifying data using multiple decision trees. And finally, RF allows assigning higher weights to minority classes during training, facilitating learning patterns in these classes.

Spore images will have a lot of internal variation, even within an image set, with some spores having larger, smaller, or out-of-focus layers. For spore segmentation, it means that data can be imbalanced, and some spore classes may be under-represented, leading to bias in the computed results. However, our proposed method resulted in a more balanced prediction with improved performance for these underclasses. Notably, this algorithm is computationally efficient and requires less computational power than other algorithms, making it well-suited for deployment in real-world applications.

Finally, it is worth noting that there are both advantages and disadvantages to the proposed method, which we will outline here.

- The use of CNN ensures that extracted features preserve pixel locality and reduce the curse of dimensionality.
- Automatic feature extraction by CNN eliminates the need to design handcrafted features, improving the method’s generalizability.
- Random forest classification enhances prediction accuracy by combining multiple decision trees and using majority voting for the final decision.
- The proposed method can be applied to solve any image-based classification and segmentation problem since CNN can learn from any image dataset.

While the proposed method has its benefits, there are also some challenges that need to be considered:

- For more complex tasks, additional features may be necessary to achieve acceptable accuracy, increasing training time and computational power.
- TEM images for the data set were obtained from two institutions. Testing the model with more data sets would be beneficial.

## Conclusion

This paper presents a novel method for spore segmentation utilizing Convolutional Neural Networks (CNN) and Random Forest (RF) decision trees. We employ multiple decision trees of a RF to enhance the classification power of the proposed method. The CNN in the proposed method employs 15 convolutional layers, ReLU layers, and 5 max-pooling layers and extracts features in TEM images and uses those features during the process of making a decision tree. The experimental results show that the method achieves good segmentation results for spores by effectively learning features. As a demonstration of the feasibility of our model, we conducted an assessment of spores that were exposed to chemical exposure. Our findings indicate the successful ability of the model to detect spores with damaged cores.

## Supporting information

**S1 Fig. The CNN algorithm automatically extracts features from images that can be used for downstream tasks. To visualize this, we passed an input image (A) through the CNN and extracted the top 8 features from layer 4 (B) and layer 10 (C), respectively. As can be seen, each layer in a CNN learns to extract different types of features from the input image and as we move deeper into the network, the features become more complex and abstract**.

**S2 Fig. Figure S2. Example images showing TEM images (left), labeled images (middle), and predicted classification (right)**.

**S3 Fig. Figure S3. Example images showing TEM images (left) and predicted classification (right) for sodium hypochlorite-treated spores**.

**S4 Fig. Figure S4. Example images showing TEM images (left) and predicted classification (right) for sodium peracetic acid-treated spores**..

## Acknowledgments

This work was supported by the Swedish Research Council (2019-04016); the Umeå University Industrial Doctoral School (IDS); Kempestiftelserna (JCK-2129.3).

The authors acknowledge the facilities and technical assistance of the Umeå Core Facility for Electron Microscopy (UCEM) at the Chemical Biological Centre (KBC), Umeå University, a part of the National Microscopy Infrastructure NMI (VR-RFI 2016-00968).

