## Supplemental information for "A hybrid CNN-Random Forest algorithm for bacterial spore segmentation and classification in TEM images"

### A step-by-step guide to install and run the program

The source code for the model implementation is available at https://github.com/sqbqamar/Spore-Segmentation

1. Open a command prompt or terminal window on your system.
2. Install the required Python libraries by running the following commands:

   i) pip install tensorflow
   ii) pip install keras
   iii) pip install opencv-python
   iv) pip install sklearn

3. Once the libraries are installed, navigate to the directory where your program file is located using the ***cd*** command.
4. Open the program file in a text editor or an IDE of your choice.
5. Load the trained model by adding the following code at the beginning of the program:

   i) from keras.models import load_model
   ii) model = load_model('path/to/your/trained/model')

Replace ***'path/to/your/trained/model'*** with the path to your trained model file.

6. Pass the input to the model by adding the following code:

   i) input_data = ## your input data
   ii) features = ## extract features from input_data using a feature extractor
   iii) numerical_data = ## convert extracted features to numerical data
   iv) prediction = model.predict(numerical_data)

Replace ***# your input data*** with the input data that you want to pass to the model. Replace ***# extract features from input_data*** with code that extracts features from the input data. Replace ***# convert extracted features to numerical data*** with code that converts the extracted features to numerical data that can be passed to the model.

7. Run the program by typing ***python file_name.py*** in the command prompt or terminal window.
8. After the program has finished running, the segmented results of using the proposed work will be displayed or saved to a file.

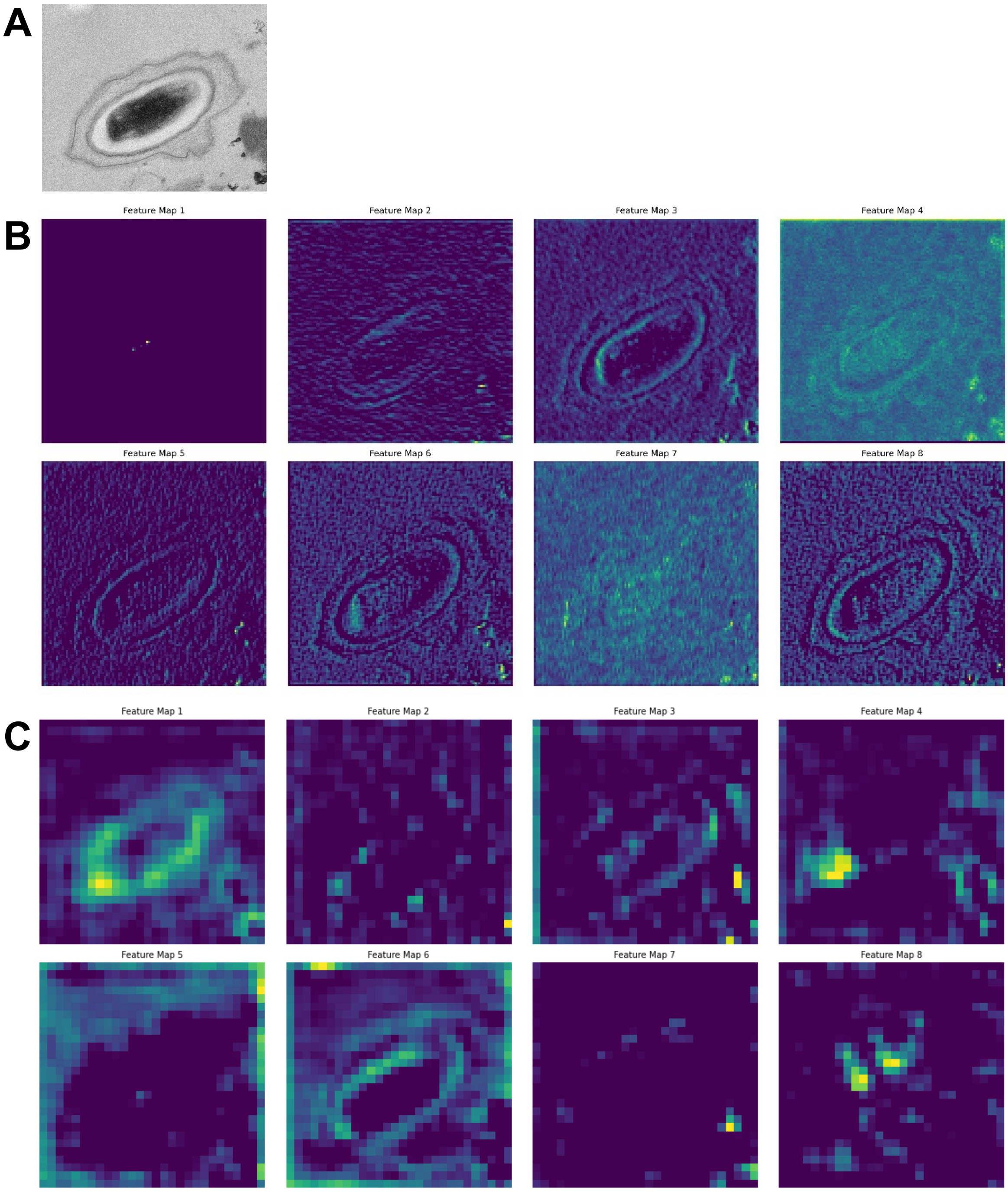


Figure S1. The CNN algorithm automatically extracts features from images that can be used for downstream tasks. To visualize this, we passed an input image **(A)** through the CNN and extracted the top 8 features from layer 4 **(B)** and layer 10 **(C)**, respectively. As can be seen, each layer in a CNN learns to extract different types of features from the input image and as we move deeper into the network, the features become more complex and abstract.

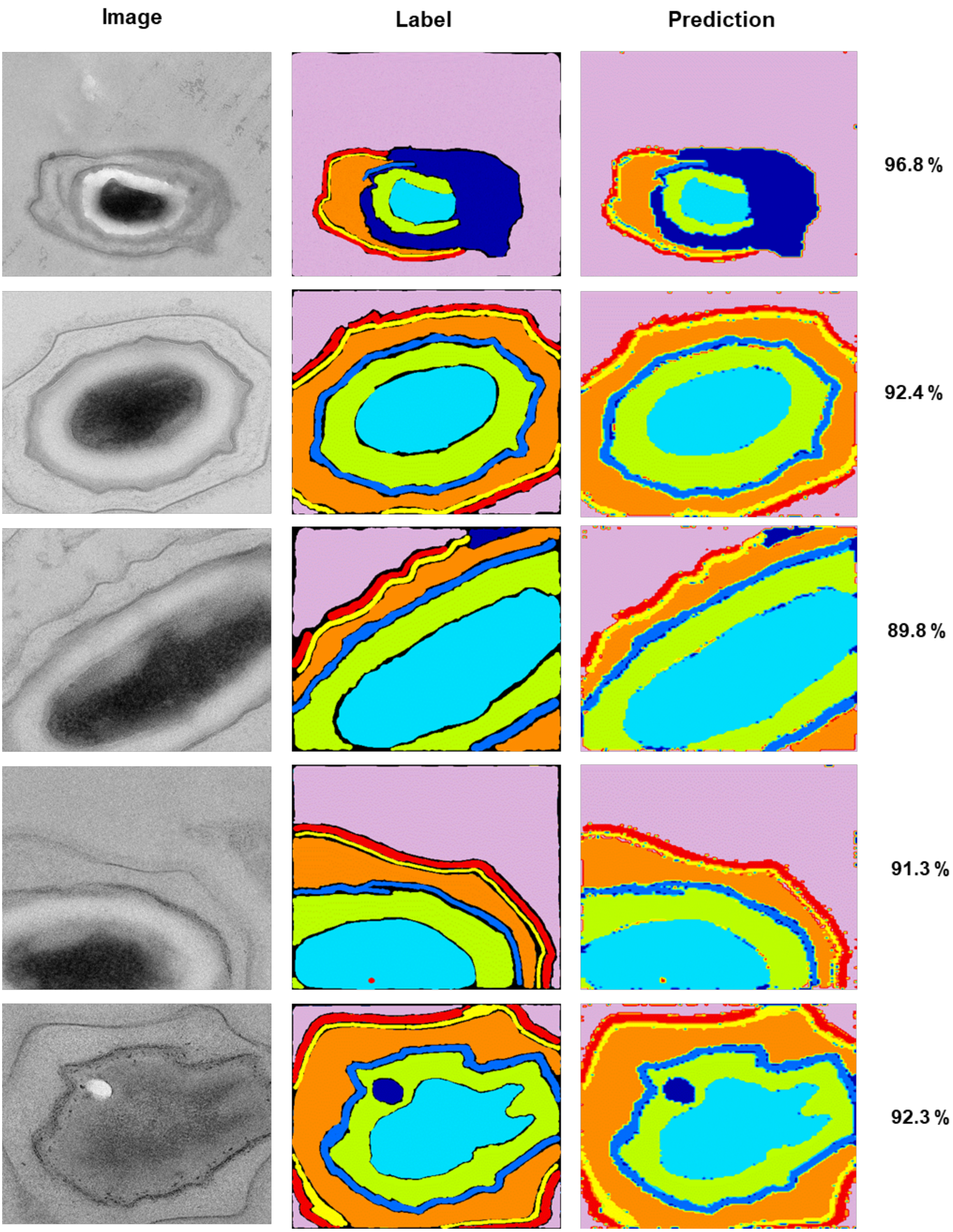
Image
Label
Prediction
96.8 %
92.4 %
89.8 %
91.3 %
92.3 %

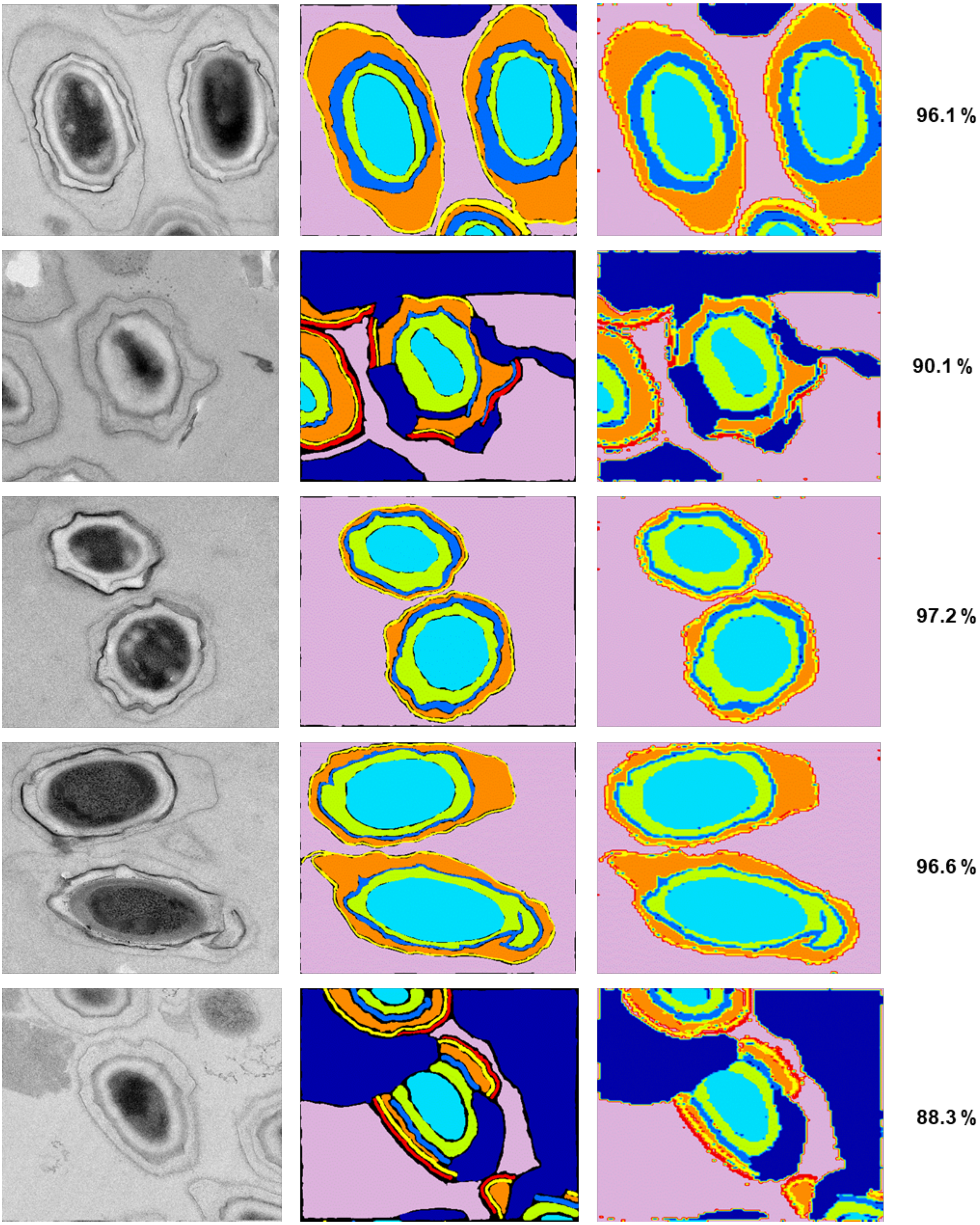


Figure S2. Example images showing TEM images (left), labeled images (middle), and predicted classification (right).

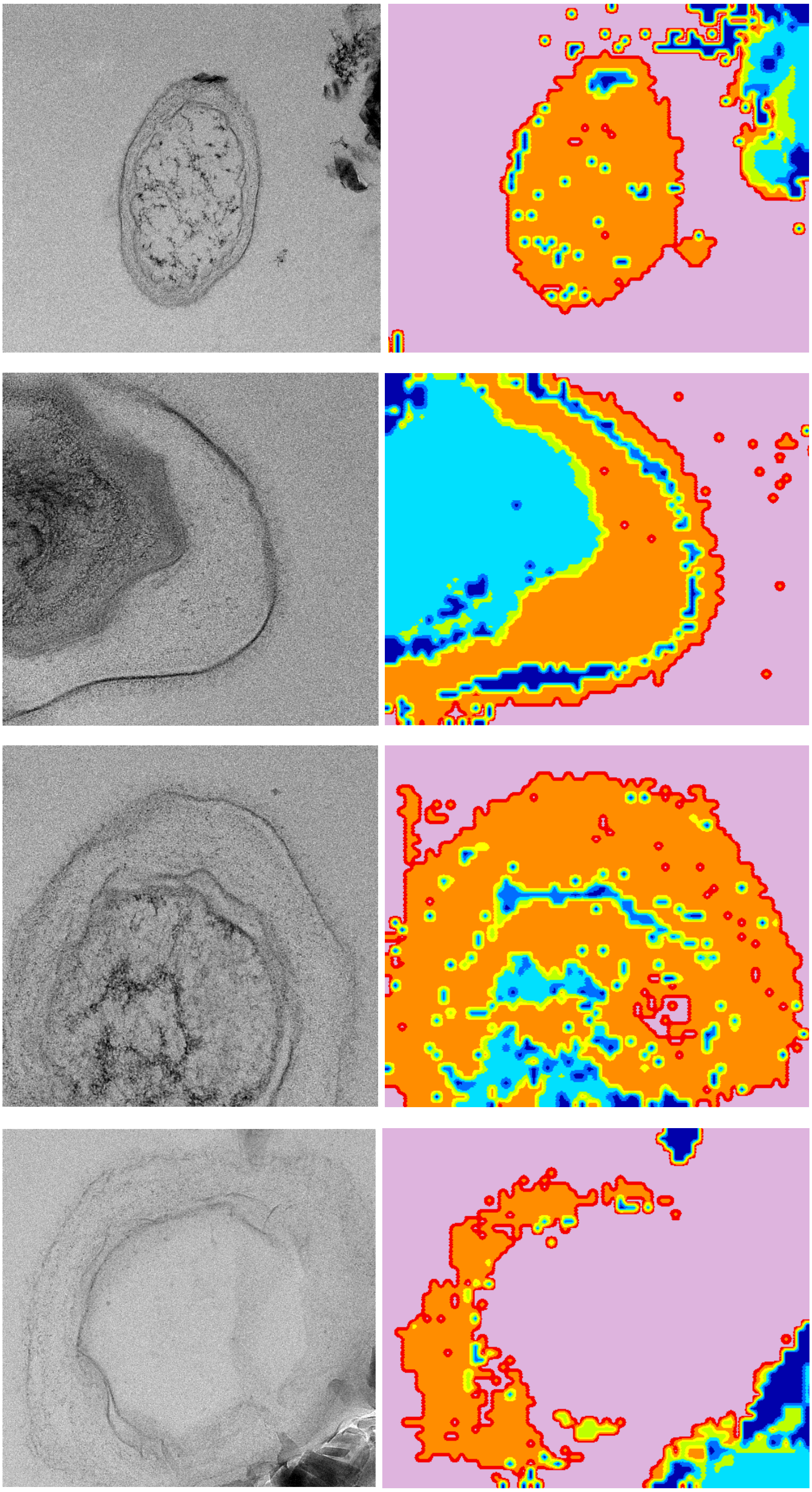

Figure S3. Example images showing TEM images (left) and predicted classification (right) for sodium hypochlorite-treated spores.

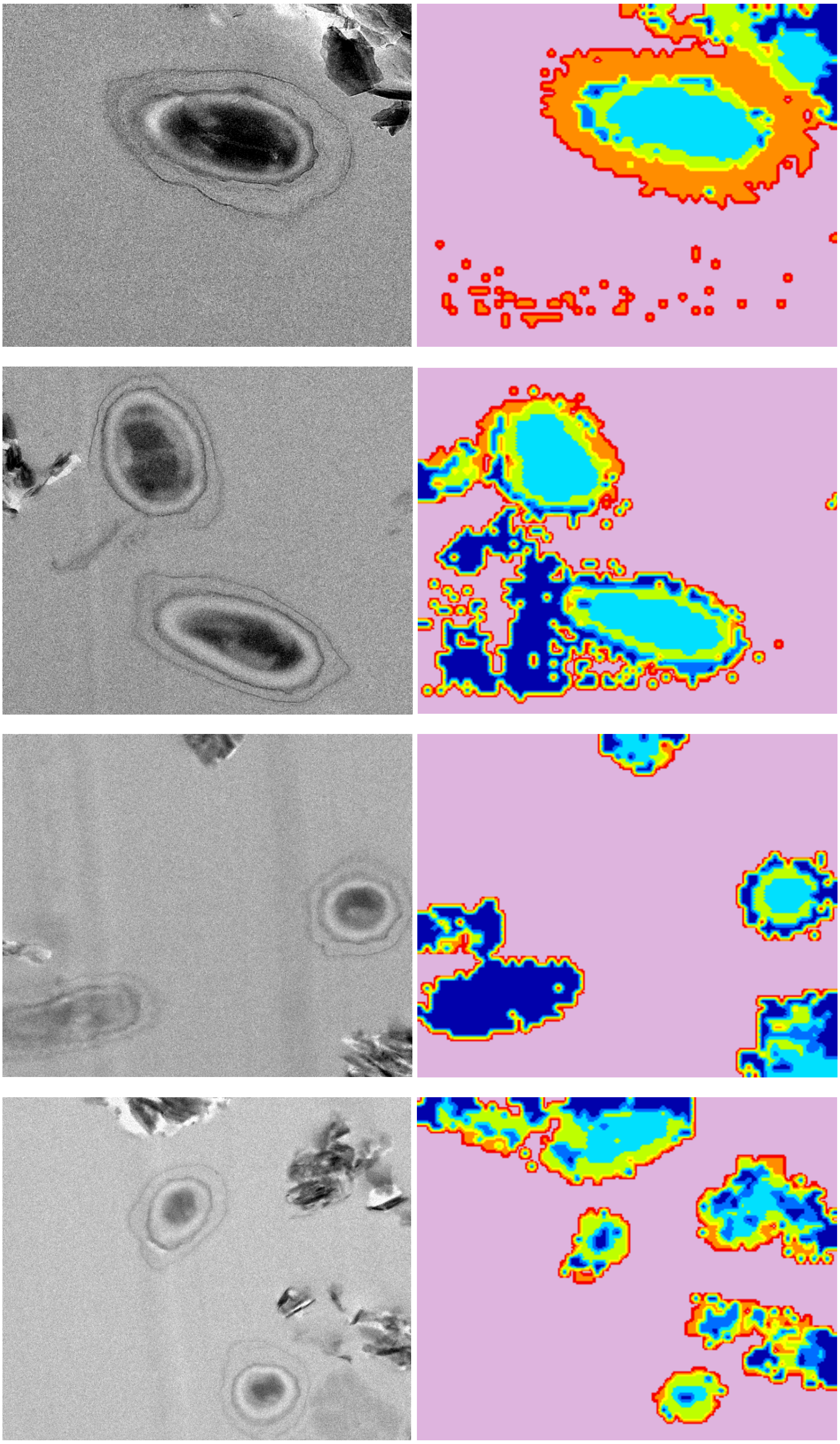

Figure S4. Example images showing TEM images (left) and predicted classification (right) for sodium peracetic acid-treated spores.
